# Simple k-RF Metrics for Comparison of Labeled DAGs

**DOI:** 10.1101/2025.07.07.663441

**Authors:** Elahe Khayatian, Louxin Zhang

## Abstract

Causal relationships between different entities are often modeled as labeled acyclic digraphs (DAGs) in biology and healthcare, in particular for depicting the progression of malignant tumor cells. Comparison of labeled DAGs is essential for developing methods for inference and evaluation of DAG models. Therefore, a robust dissimilarity metric is critical for such comparison tasks. We introduce new dissimilarity measures for labeled DAGs by refining the k-Robinson-Foulds distance, originally defined to compare labeled trees. The new measures are defined based on the comparison of local node-induced multisets of labels. They can be used to compare DAGs with different label sets, without the need to introduce auxiliary nodes or remove existing ones.

## 1 Introduction

NEtworks (or graphs) are widely used as mathematical models to represent a binary relation between entities, such as individuals in social networks [27], proteins and drugs in biological networks [7], and species in evolutionary networks [13]. Network modeling often provides valuable insights into these relationships. For example, drug interaction networks help in prediction of the synergistic effects of drug combinations [7]. Phone call networks facilitate the discovery of communication patterns within a community [27]. Evolutionary networks help uncover the evolutionary history of species [13].

Since these binary relations are often asymmetric, acyclic digraphs (DAGs) are commonly used for modeling. For instance, DAGs find applications in epidemiology [2] and pediatrics [34]. DAGs are also used to design deep learning methods for action recognition tasks ([31], [33]) and generating gene expression data ([12], [36]).

Consequently, researchers are particularly motivated to propose metrics for effectively assessing similarity/dissimilarity between DAGs. These dissimilarity measures play important roles in DAG aggregation [23] and learning, having wide applications in biology and health-care [35].

A number of dissimilarity metrics have been proposed in the space of phylogenetic networks ([3], [4], [5], [6], [15], [17], [21], [22], [25], [32]). Since phylogenetic networks have only leaves labeled, these dissimilarity measures are not ideal to compare the DAGs in which all nodes are labeled [11], [19], which also appear frequently in other fields. For example, in a pediatric study [34], casual relationship is represented as a DAG in Figure 1, where each node has a label.

**Fig. 1.**
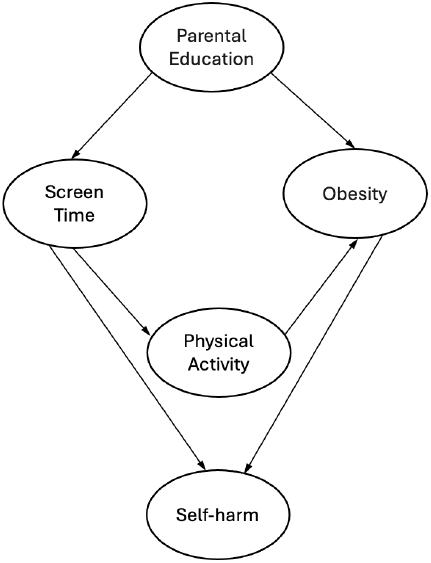
A labeled DAG (directed acyclic graph) model in a pediatric study (redrawn from [34]). It illustrates the following causal relationships: Low parental education leads to both increased screen time (e.g., watching TV, playing video games) and a higher risk of obesity in children. Increased screen time directly reduces physical activity, which further raises the risk of obesity. Both increased screen time and obesity have also been associated with greater risks of low self-esteem and self-harm in adolescents.

In cancer biology, labeled DAGs are used to model the progression trajectory of tumor evolution [9], [10], [20], [24], [29], providing valuable information for cancer diagnosis and treatment [10]. In these models, each node represents a cell population and is labeled with a set of mutations that characterize the population. Metrics for comparing cancer progression models (CPMs) have been proposed in [11], [16], and [19].

Katz Graph Similarity (KGS) [26] is another kind of dissimilarity metric proposed for various applications of DAGs. When comparing two DAGs, KGS first computes the Katz Similarity Vector (KSV) for each DAG [26] to capture node similarity, and then the resulting KSVs are compared. A related measure is the Generalized Kendall-tau [23] that quantifies similarity based on the pairwise dissimilarity between corresponding node pairs in the two DAGs. Both KGS and Generalized Kendall-tau are parameter-based similarity measures.

The above-mentioned metrics share a common weakness: they are not efficient for comparing DAGs with different label sets. For example, Malmi et al. [23] suggest adding isolated nodes to align label sets, while Nayak et al. remove differing labels in their simulation experiments in [26]. However, these strategies may alter the structure of the input DAGs or artificially increase the measured dissimilarity between them.

In this work, we propose a new class of dissimilarity measures for labeled DAGs. They are developed from the *k*-RF distance proposed for labeled trees in [19] and are thus called the simple and modified *k*-RF measures. The new measures are defined based on the comparison of local node-induced multisets of labels.

These measures have been shown to be both metric and computationally efficient in the space of 1-labeled DAGs. Moreover, they can be naturally extended to compare DAGs with different label sets, without the need to introduce auxiliary nodes or remove existing ones.

Our contributions to labeled DAG comparison are presented in the next eight sections. In Section 2, basic concepts and notation that are used in this study are introduced. In Section 3, we define the simple *k*-RF distances for rooted 1-labeled trees. In Section 4, the measures defined in Section 3 are generalized for 1-labeled DAGs. Their mathematical properties are investigated. In Section 5, considering the limitation of the simple *k*-RF distances, the modified *k*-RF measures are introduced. In Section 6, the simple and modified *k*-RF measures are compared with some existing measures. In Section 7, the simple *k*-RF and the *k*-RF measures are compared for trees with different label sets. Finally, we conclude the paper by summarizing the key points in Section 8.

## 2 Concepts and Notation

### 2.1 Graphs and Digraphs

A *graph* or *digraph G* is a mathematical structure consisting of a set of nodes *V* (*G*) and a set of edges *E*(*G*). In a graph, each edge is an unordered pair of nodes. If *u* and *v* are joined by an edge *e* in a graph, we can write *e* = (*u, v*) or *e* = (*v, u*). In a digraph, each edge is an ordered pair of nodes.

Let *G* be a graph or digraph. If *e* = (*u, v*) ∈ *E*(*G*), *u* and *v* are said to be *adjacent* to each other; *e* is incident to *u* and *v*; and *u* and *v* are the endpoints of *u*. The degree *d*(*u*) of a node *u* is defined as the number of edges incident to it.

Furthermore, in a digraph *G*, if *e* = (*u, v*) ∈ *E*(*G*), *u* and *v* are called the *tail* and *head* of *e*, respectively; *e* is called an outgoing edge of *u* and an incoming edge of *v*. The *indegree* and *outdegree* of a node *u* are defined as the number of its incoming and outgoing edges, respectively, which are denoted as *d*_*in*_(*u*) and *d*_*out*_(*u*). Clearly, the degree of *u* is equal to the sum of the indegree and outdegree of *u*.

In a graph, the nodes of degree 1 are called leaves. In a digraph, the nodes of outdegree 0 are called leaves. The set of all leaves of a graph or dirgraph *G* is denoted by Leaf(*G*). The nodes of indegree 0 are called the sources. Non-leaf nodes are called internal nodes.

A *path* of length *k* from *u* to *v* in a graph or digraph *G* is a sequence of nodes *u* = *u*_0_, *u*_1_, *u*_2_, …, *u*_*k*_ = *v* such that (*u*_*i*_, *u*_*i*+1_) ∈ *E*(*G*) for any 0 ≤ *i* ≤ *k* − 1. A path from *u* to itself is called a *cycle*.

The *distance* from *u* to *v* in a graph or digraph *G* is the length of the shortest paths from *u* to *v*, denoted as *d*_*G*_(*u, v*) or simply *d*(*u, v*). We set *d*_*G*_(*u, v*) = ∞ if there is no path from *u* to *v* in *G*.

We define the diameter of a graph *G* as max_*u,v* ∈ *V* (*G*)_ {*d*_*G*_(*u, v*)}. For a digraph *G*, the diameter is defined as the diameter of the graph with the node set *V* (*G*) and the edge set *E*(*G*), where the order on each *e* ∈ *E*(*G*) is ignored.

A graph *T* is called a *tree* if there exists exactly one path from any *u ∈ V* (*T*) to any *v ∈ V* (*T*).

An edge (*u, v*) *∈ E*(*G*) is said to be *redundant* if there is a path with length *k* ≥ 2 from *u* to *v*.

### 2.2 Labeled Graphs and Digraphs

Let *X* be a set and ℙ (*X*) denote the set of all subsets of *X*. A graph or digraph *G* is labeled over *X* if it is equipped with a function *ℓ* : *V* (*G*) → ℙ (*X*)*\*{∅} such that ∪_*v* ∈ *V* (*G*)_*ℓ*(*v*) = *X*. In particular, if each *ℓ*(*v*) has only one element and *ℓ* is one-to-one, *G* is said 1*-labeled* over *X*, where we identify each node *v* with its label *ℓ*(*v*).

Two 1-labeled graphs or digraphs *G* and 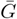 are *identical* if and only if 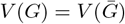 and 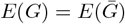.

In this paper, a labeled graph or digraph is assumed to be 1-labeled.

### 2.3 DAGs and Rooted Trees

A digraph *G* is called a *DAG* if it contains no cycle. It is called a *rooted DAG* if it has exactly one source. Let *G* be a rooted DAG and *u, v* ∈*V* (*G*) be distinct. If (*u, v*) ∈ *E*(*G*), *u* is said to be a *parent* of *v* and *v* a *child* of *u*. More generally, *u* is called an *ancestor* of *v* and *v* is called a *descendant* of *u* if there is a path from *u* to *v* with length at least one. The sets of the ancestors and descendants of *u* are denoted by *A*_*G*_(*u*) and *D*_*G*_(*u*), respectively.

The *level* of a node *w* ∈ *V* (*G*) is defined to be the maximum length of a directed path from a source to *w*. The depth(*G*) is the maximum level of a node.

A rooted tree is a rooted DAG in which every non-root node has exactly one parent.

### 2.4 Multisets and Their Operations

A collection of elements is called a multiset if each element can occur a finite number of times. If *A* is a multiset and *x* ∈ *A*, the number of occurrences of *x* in *A* is denoted as *m*(*x, A*). Thus, *A* can be represented as *A* = {*x*^*m*(*x,A*)^ : *x* ∈ Supp(*A*)}, where Supp(*A*) denotes the set of all distinct elements in *A*.

Let *A* and *B* be two multisets. We say *A* ⊆_*m*_ *B* if and only if Supp(*A*) ⊆ Supp(*B*) and ≤ *m*(*x, A*) ≤ *m*(*x, B*) for any *x* ∈ Supp(*A*).

We also say *A* = *B* if and only if Supp(*A*) = Supp(*B*) and *m*(*x, A*) = *m*(*x, B*) for any *x ∈* Supp(*A*).

In addition, we also define the following operations on multisets:

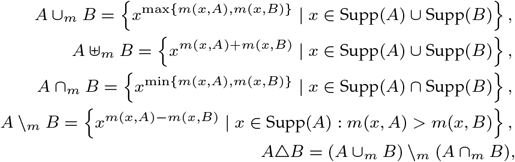

where *m*(*x, X*) = 0 if *x ∈/ X* for *X* = *A, B*.

### 2.5 Multi-labeled DAGs

Let *X* be a set and ℳ (*X*) denote the collection of multisets with Supp(*X*). A DAG *G* is called a *multi-labeled DAG* on *X* if *G* is equipped with a function *ℓ* : *V* (*G*) *→* ℳ (*X*) such that *∪*_*v∈V* (*G*)_Supp(*ℓ*(*v*)) = *X* and *ℓ*(*v*)*≠ ∅* for every *v ∈ V* (*T*).

### 2.6 Resolution of Dissimilarity Measures

let D be a space of labeled (multi-labeled) DAGs and *d, d*^*′*^ : 𝒟 *×* 𝒟 *→* [0, 1] be two dissimilarity measures. We say that *d* has a higher resolution than *d*^*′*^ on 𝒟 if |*d*(𝒟 *×* 𝒟)| *>* |*d*^*′*^(𝒟 *×* 𝒟)|, where

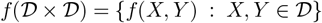

for *f* = *d, d*^*′*^.

## 3 Simple *k*-RF Distances for Labeled Trees

The authors use the *k*-neighborhood to define the *k*-RF metrics in the space of rooted labeled trees [19].

Let *T* = (*V, E*) be a rooted labeled tree and let *k* ≥ 0. Each edge *e* = (*u, v*) ∈ *E* induces an ordered pair *P*_*T,k*_(*e*) = (*D*_*T,k*_(*v*), *B*_*T,k*_(*u*) *\ D*_*T,k*_(*v*)) of label subsets, where *D*_*T,k*_(*v*) and *B*_*T,k*_(*u*) are respectively defined as:

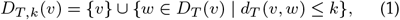

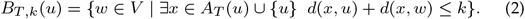

Additionally, define 𝒫 (*T, k*) = {*P*_*T,k*_(*e*) | *e ∈ E*(*T*)}.

### Definition 1.

([19]) *Let X and Y be two rooted labeled trees. The k-RF distance between X and Y is defined as:*

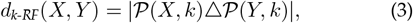

*where Δ is the symmetric set difference operator*.

In the rest of this section, we will simplify the metric to develop a computationally efficient measure for labeled DAGs.

### 3.1 Definition

Let *T* be a rooted labeled tree and let *k* ≥ 0. We define

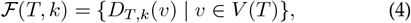

where *D*_*T,k*_(*v*) is defined in Eqn (1) and is called *k*-descendant-hood of *v* in *T*.

#### Definition 2.

*Let X and Y be two rooted labeled trees. The simple k-RF (s-k-RF) dissimilarity measure between X and Y is defined as follows*.

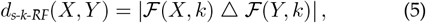

*where Δ is the symbol for symmetric set difference*.

#### Observation 1.

*Let T be a rooted labeled tree and k* ≥ 0. *In addition, let r be the root of T*. *Then, D*_*T,k*_(*r*) = *B*_*T,k*_(*r*).

#### Observation 2.

*Let T be a rooted labeled tree and k* ≥ 0. *In addition, assume v ∈ V* (*T*) *and* |*V* (*T*)| *>* 1. *Then, we have:*

- *If v is a non-root internal node, there exist two nodes u, w* ∈*V* (*T*) *such that* (*u, v*), (*v, w*) ∈ *E*(*T*). *As such, computing both D*_*T,k*_(*v*) *and B*_*T,k*_(*v*) *is necessary to calculate P*_*T,k*_(*e*_1_) *and P*_*T,k*_(*e*_2_), *where e*_1_ = (*u, v*) *and e*_2_ = (*v, w*).
- *If v is the root of T, there is at least one node x* ∈ *V* (*T*) *such that e* = (*v, x*) ∈ *E*(*T*). *Thus, by Observation 1, computing D*_*T,k*_(*v*) *is needed to compute P*_*T,k*_(*e*).
- *If v is a leaf of T, there is at least one node y* ∈ *V* (*T*) *such that e* = (*y, v*) ∈ *E*(*T*). *Thus, computing D*_*T,k*_(*v*) *is needed to obtain P*_*T,k*_(*e*).

Observation 2 highlights that for a rooted labeled tree *T* with *n* nodes, computing all *n* sets in ℱ (*T, k*) is required to calculate 𝒫 (*T, k*); additionally, computing *B*_*T,k*_(*v*) for each non-root internal node *v* ∈ *V* (*T*) is needed to obtain 𝒫 (*T, k*). Furthermore, *n* − 1 set difference operations are needed to obtain the second term of ordered pairs in 𝒫 (*T, k*). As such, the s-*k*-RF distances could enhance the computation efficiency and space complexity of the *k*-RF distances, making *d*_s-k-RF_ a simplified version of *d*_k-RF_. We now explore the relation of the s-*k*-RF distances with the traditional RF distance.

#### Definition 3.

([28]) Let *X* and *Y* be two rooted labeled trees. The RF distance between *X* and *Y* is defined as:

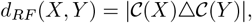

where 𝒞 (*T*) = {*D*_*T*_ (*v*) *∪* {*v*} : *v ∈ V* (*T*)} for *T* = *X, Y*.

#### Proposition 1.

*Let S and T be two rooted labeled trees and m* = max{*depth*(*S*), *depth*(*T*)}. *The s-k-RF distance between S and T is equal to d*_*RF*_ (*S, T*) *for any k* ≥ *m*.

**Proof**. Note that if *k* ≥ *m, D*_*X,k*_(*v*) = *D*_*X*_ (*v*) ∪ {*v*} for any *v* ∈*V* (*X*) in *X* = *S, T*. The result is derived from Definitions 2 and 3.

### 3.2 Mathematical Properties

#### Lemma 1.

*For every three sets A, B, S*,

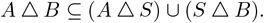

**Proof**. Let *x ∈ A Δ B* = (*A \ B*) *∪* (*B \ A*). Without loss generality, we may assume *x ∈ A \ B*. Then, if *x ∈ S, x ∈ S \ B ⊆ S Δ B*. If *x ∈/ S*, then *x ∈ A \ S ⊆ A Δ S*.

#### Lemma 2.

*Let T be a rooted labeled tree* such *that* |*V* (*T*) | ≥ 2 *and L be a subset of Leaf* (*T*). *Define T* ^*′*^ *to be the tree obtained by deleting the leaves of L together with the edges incident to these leaves. Then, for any k* ≥ 1,

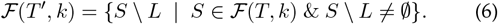

**Proof**. Note that *V* (*T* ^*′*^) = *V* (*T*) *\ L*. For each *v ∈ V* (*T* ^*′*^), by definition, 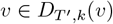 and thus 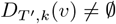.

Let *u ∈ V* (*T* ^*′*^). If 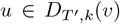, then 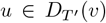 and 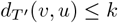. Since any directed path of *T* ^*′*^ is also a path in 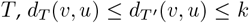. This implies that *u* ∈ *D*_*T,k*_(*v*). Since *u* ∉ ℒ, *u* ∈ *D*_*T,k*_(*v*) \*L*. Thus,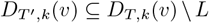.

Conversely, for any *u* ∈ *D*_*T,k*_(*v*)*\L, u* ∉ *L* and there is a directed path *P* from *v* to *u* with length at most *k*. Since a leaf cannot appear in the middle of a path, the path *P* is also a path in *T* ^*′*^. This implies that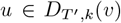. Thus,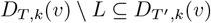.

Taken together, the above two facts imply that 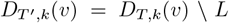 for each *v* ∈ *V* (*T* ^*′*^). Therefore, Eqn. (6) holds.

#### Theorem 2.

*Let k* ≥ 1 *be an integer. The s-k-RF defined in Eqn. (5) is a metric in the space of rooted labeled trees*.

**Proof**. The non-negativity and symmetry properties can be observed from the definition of the dissimilarity measure.

Let *X, Y, Z* be three rooted labeled trees. Setting *A* = ℱ (*X, k*), *B* = ℱ (*Y, k*) and *S* = ℱ (*Z, k*), we obtain:

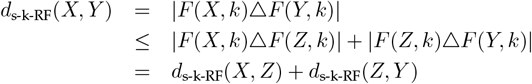

from Lemma 1. This proves the triangle inequality.

Lastly, we prove that *d*_s-k-RF_(*X, Y*) = 0 implies that *X* and *Y* are identical for any two rooted labeled trees *X* and *Y*. Clearly. the value 0 of the measure implies that *V* (*X*) = *V* (*Y*). For the purpose, we just need to prove that *E*(*T*) can be uniquely determined by ℱ (*T, k*) using the mathematical induction on |*V* (*T*) |.

If *T* consists of a singe node *v*, then ℱ (*T, k*) = {*v*}. If *T* has two or more nodes, the root *r* of *T* has at least one child. Since *k* ≥ 1, *D*_*T,k*_(*r*) contains at least two nodes. For any leaf *ℓ* of *T, D*_*T,k*_(*ℓ*) = {*ℓ*}. This implies that (*T, k*) contains at least 2 distinct *k*-descendant sets. Therefore, *T* is the single-node tree if and only if ℱ (*T, k*) is a singleton.

Assume *E*(*M*) is uniquely determined by ℱ (*M, k*) for arbitrary rooted labeled tree *M* such that |*V* (*M*) | *< t*. We consider a rooted labeled tree *T* such that |*V* (*T*) | = *t*.

There is no edges leaving a leaf. Since *k* ≥ 1, *D*_*T,k*_(*v*) = {*v*} if and only if *v* is a leaf. Therefore, we can identify all leaves of *T* from the singletons of (*T, k*).

For *v* ∈ *V* (*T*) *\ Leaf* (*T*), there is a unique edge *e* = (*u, v*) entering any *v*. Since *k* ≥ 1, the children of *v* are all leaves if and only if *D*_*K*_(*v*) \*Leaf* (*T*) = {*v*}. Therefore, we can identify *v* whose children are all leaves from the subsets *S* ∈ ℱ (*T*) such that *S* \ *Leaf* (*T*) contains only *v*.

Let *V* ^*′*^ be the set of all nodes whose children are just leaves and 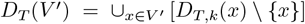. Since *V* ^*′*^ is nonempty, *D*_*T*_ (*V* ^*′*^)*≠ ∅*. Define *E*^*′*^(*T*) = {(*x, y*) ∈ *E*(*T*) | *x ∈ V ′, y ∈, D*_*T*_ (*V* ^*′*^)}.

For the tree *T*^*′*^ obtained from *T* by the removal of the leaves in *D*_*T*_ (*V* ^*′*^), |*V* (*T* ^*′*^)| = |*V* (*S*) | −|*D*_*T*_ (*V* ^*′*^)| *< k*. By Lemma 2, ℱ (*T* ^*′*^, *k*) can be efficiently constructed from ℱ (*T, k*). By the induction hypothesis, *E*(*T* ^*′*^) is uniquely determined by ℱ (*T* ^*′*^, *k*). As a result, *E*(*T*) = *E*(*T* ^*′*^) ∪ *E*^*′*^(*T*) is determined. This concludes the proof.

### 3.3 Frequency Distribution of S-*k*-RF Distances

The simulation experiments reported in [19] suggest that the distribution of the *k*-RF distances between labeled trees varies with different values of *k*. This observation was recently confirmed by Fuchs and Steel in [14]. They mathematically proved that a linear transformation of (*n* − 2)-RF distance follows a normal distribution, whereas that of 0-RF distance has a Poisson distribution.

Here, we investigate whether the untransformed s-*k*-RF distance in the space of rooted labeled trees follows a normal or Poisson distribution.

To this end, we computed pairwise s-*k*-RF distances between labeled *n*-node rooted trees with the same label set for *n* = 4, 5, 6. The resulting histograms are given in Figures 2, 3, 4, respectively, where the Gaussian fits were obtained using the following `curve_fit` function

**Fig. 2.**
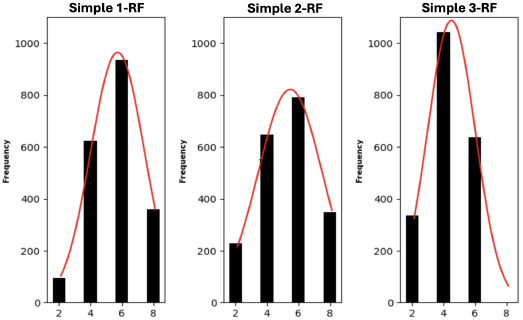
The distributions of s-*k*-RF scores for *k* = 1, 2, 3 in the space of rooted labeled 4-node trees with the same label set. The red curve is a Gaussian fit.

**Fig. 3.**
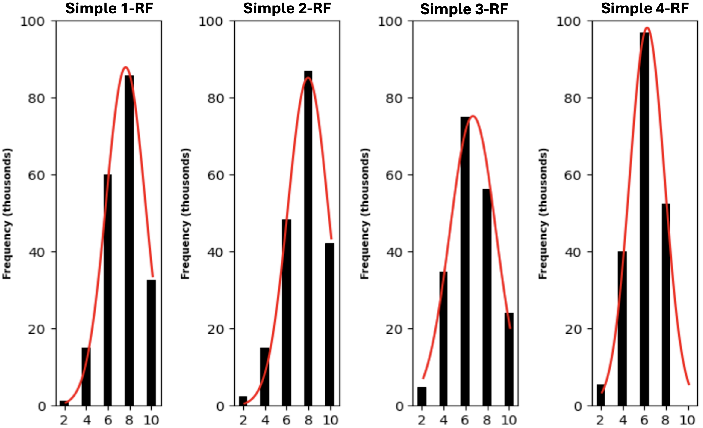
The distribution of s-*k*-RF scores for *k* = 1, 2, 3, 4 in the space of rooted labeled 5-node trees with the same label set. The red curve is a Gaussian fit.

**Fig. 4.**
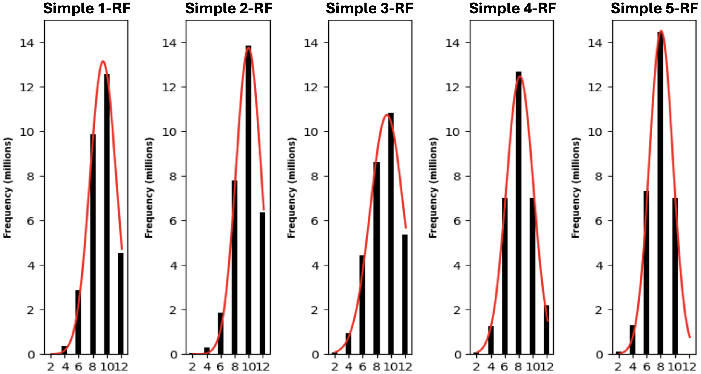
The distributions of s-*k*-RF distances in the space of rooted labeled 6-node trees with the same label set for *k* = 1, 2, 3, 4, 5. The red curve is a Gaussian fit.

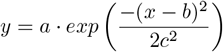

from the SciPy library, which approximates a normal distribution. In Section IV of Supplementary Document, Figures S1, S2, and S3 show the distribution of pairwise s-*k*-RF distances between labeled *n*-node rooted trees with the same label set for *n* = 7, 10, 13, respectively.

These results suggest that the distribution of the s-*k*-RF distances appears to be normal for each *n* and for all *k* = 1, …, *n −* 1.

### 3.4 Correlation of S-*k*-RF Distances with Several Distances

Using a Pearson correlation analysis, we compared different s-*k*-RF distances with several existing distances.

We first generated a set of random trees with 30 nodes using a program described in [16]. In each generated tree, the root was always labeled 0, while the remaining vertices were labeled with integers from 1 to 29.

After creating the dataset, we sampled 950 labeled rooted trees from the dataset. We divided the trees into 5 families and changed the labels of some nodes of the trees within the last four families so that trees in each family shared the same label set and trees from different families had distinct but overlapping label sets.

We then computed all pairwise distances between trees within each family, using s-*k*-RF scores for 1 ≤ *k* ≤ 21; CASet ∩ [11]; DISC ∩ [11]; and GRF [22] (Section I, Supplementary Document). Finally, we computed Pearson correlation of each s-*k*-RF measure with the other three measures. As illustrated in the left plot of Figure 5, we observed the following facts.

**Fig. 5.**
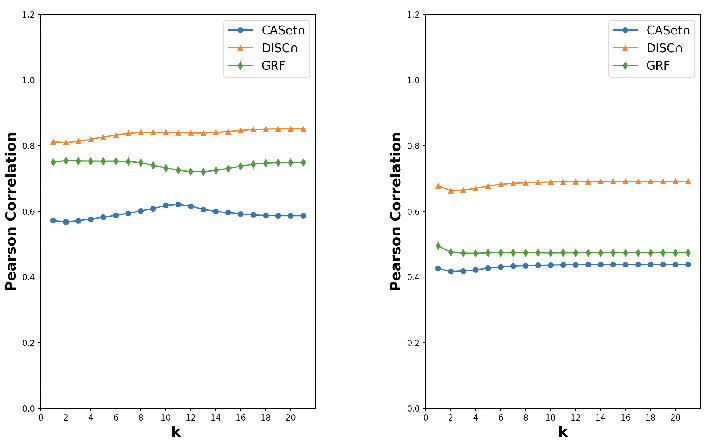
Pearson correlation of s-*k*-RF measures with three existing measures CASet∩ [11], DISC∩ [11], and GRF [22]. The analysis was conducted on random trees with the same label sets (left) and on trees with different but overlapping label sets (right).

- Each s-*k*-RF measure indicated a strong correlation with DISC∩ and less correlation with CASet∩.
- There was no clear change in the correlation values when *k* increased from 1 to 21.
- The correlation values were between 0.55 and 0.85.

Next, we computed the distance for any pair containing one tree from the first family and one tree from other families, using each of the above-mentioned dissimilarity measures. Then, we computed Pearson correlation of each s-*k*-RF measure with each of the other three measures. In this case, we observed the following in Figure 5 (right).

- Each s-*k*-RF measure indicated a higher correlation with DISC∩ and less correlation with CASet∩.
- The correlation values were between 0.4 and 0.7.

As Figure 5 shows, the correlation between each s-*k*-RF (1≤ *k* ≤ 21) and each of the above three measures was higher in the case where each investigated pair contained trees with the same label set.

### 3.5 Correlation of S-*k*-RF and *k*-RF Distances

We further analyzed the Pearson correlation between the s-*k*-RF distance and the *k*-RF distance.

For this purpose, we utilized the dataset described in Section 3.4. Recall that the dataset contained 5 tree families; additionally, trees from each family shared the same label set, whereas the trees across the families had different but overlapping label sets.

We then computed the *k*-RF (1≤ *k* ≤ 21) distance between any two trees from each family and between any two trees *T* and *S*, where *T* was from the first family and *S* was from the other families.

The Pearson correlation coefficients between the s-*k*-RF and the *k*-RF (1 ≤ *k* ≤ 21) are presented in Figure 6. The analyses suggest:

**Fig. 6.**
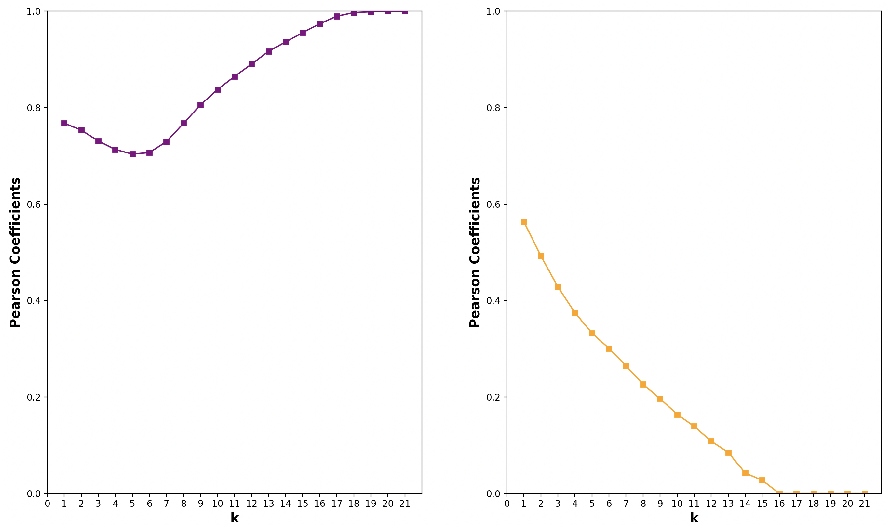
Pearson correlation of the s-*k*-RF measure with the *k*-RF measure (1≤ *k* ≤ 21). The analysis was conducted on trees with the same label sets (left) and on trees with different but overlapping label sets (right).

- For each *k*, the s-*k*-RF and the *k*-RF distance were positively correlated.
- The correlation values were higher when the distance values between trees with the same label set were considered.
- When *k* increased from 6 to 21, the correlation values increased when distances between trees with the same label set were used. In contrast, the values dropped when distances between trees with different but overlapping label sets were used.

## 4 S-*k*-RF Distances for Labeled DAGs

Let *k* ≥ 0. Clearly, *D*_*G,k*_(*v*) in Eqn. (1), ℱ (*X, k*) in Eqn. (4) are well defined for DAGs. Consequently, the s-*k*-RF distance given in Definition 2 naturally extends to DAGs. However, this dissimilarity measure is no longer a metric in the space of DAGs. For instance, for the two distinct labeled DAGs *G* and 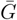 in Figure 7, we have 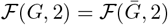 and thus their s-2-RF score is 0.

**Fig. 7.**
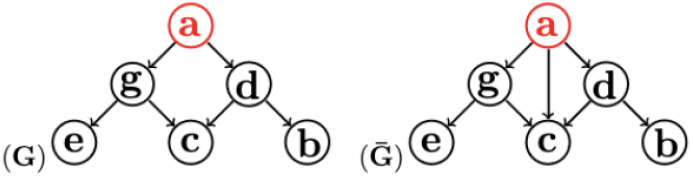
Two rooted labeled DAGs whose refined s-2-RF score is 2.

To address this issue, we redefine the s-*k*-RF distance by incorporating the number of paths of certain lengths between pairs of nodes into the definition.

For each *u* ∈ *D*_*G,k*_(*v*) (defined in Eqn. (1)), we define the *k*-reach multiplicity *n*(*v, u*) of *u* as the number of directed paths from *v* to *u* with length at most *k*. We then redefine the *k*-descendant-hood of *v* as:

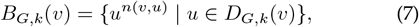

and

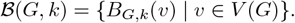

### Definition 4.

*For two labeled DAGs G and* 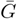, *the s-k-RF distance between G and* 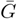 *is defined as:*

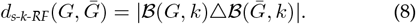

### Example 1.

*Consider the rooted labeled DAGs given in Figure*

*7. The sets of 2-descendant-hoods associated with all the nodes in the two DAGs are listed below*.

| $v$ | $B_{G,2}(v)$ | $B_{\bar{G},2}(v)$ |
| --- | --- | --- |
| $a$ | $\{a, b, c^2, d, e, g\}$ | $\{a, b, c^3, d, e, g\}$ |
| $b$ | $\{b\}$ | $\{b\}$ |
| $c$ | $\{c\}$ | $\{c\}$ |
| $d$ | $\{b, c, d\}$ | $\{b, c, d\}$ |
| $e$ | $\{e\}$ | $\{e\}$ |
| $g$ | $\{e, c, g\}$ | $\{e, c, g\}$ |

*Therefore*, 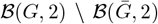 *contains only* {*a, b, c*^2^, *d, e, g*}, *whereas* 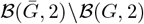 *contains only* {*a, b, c*^3^, *d, e, g*}. *Therefore, the s-*2*-RF distance between G and* 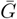 *is* 2.

### 4.1 Mathematical Properties

#### Lemma 3.

*The s-k-RF in Eqn*.*8 satisfies the non-negativity, symmetry and triangle inequality conditions*.

**Proof**. The non-negativity and symmetry conditions are trivial. Setting *A* = ℬ (*G*_1_, *k*), *B* = ℬ (*G*_2_, *k*), and *S* = ℬ (*G*_3_, *k*), by Lemma 1, we obtain

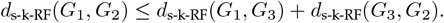

#### Observation 3.

*Let G be a labeled DAG and x, y* ∈ *V* (*G*) *such that x* ≠ *y. Then, if p is a directed path from x to y in G, the level of y in G is greater than the level of x in G*.

#### Lemma 4.

*Let G be a DAG and let S consist of the source nodes in G. For any X* ⊆ *S, G*_*X*_ *denotes the DAG obtained from G by removing all nodes in X and all the edges incident to the nodes in X. Then, for each X ⊆ S and each k* ≥ 1,

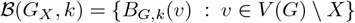

**Proof**. The proof follows from the fact that a source node has an indegree of 0 and thus is not a descendant of any other node.

#### Theorem 3.

*Let k* ≥ 1 *be an integer. The s-k-RF is a metric in the space of all labeled DAGs*.

**Proof**. By Lemma 3, we just need to show that 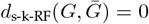 implies that *G* and 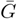 are identical for any two labeled DAGs *G* and 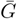. First, 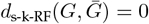 implies that 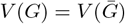.

To prove the statement, we show that each *V*_*l*_(*G*) = {*v* ∈ *V* (*G*) | the level of v is *l*} for 0 *≤ l ≤* depth(*G*) and *E*(*G*) are uniquely determined by ℬ (*G, k*) using mathematical induction on *n* = |*V* (*G*)|.

Note that as *k* ≥ 1, *s* is a source if and only if *B*_*G,k*_(*s*) is the only multiset in ℬ (*G, k*) that contains *s*. Therefore, each source of *G* is uniquely determined by ℬ (*G, k*).

Now, assume *n* = 1. Then, *V* (*G*) = {*v*} is a singleton. As *G* is a DAG, *v* is a source, *E*(*G*) = ∅ and depth(*G*) = 0. Thus, *V*_0_(*G*) = {*v*} and *E*(*G*) are uniquely determined by *B*(*G, k*) = {{*v*}}.

Note that if *n* ≥ 1, *E*(*G*) = ∅ if and only if all nodes in *V* (*G*) are sources, if and only if for any *v* ∈*V* (*G*), *B*_*G,k*_(*v*) = {*v*}. Therefore, if *E*(*G*) = ∅, then depth(*G*) = 0. Additionally, *V*_0_(*G*) = *V* (*G*) and *E*(*G*) are uniquely determined by *B*(*G, k*). Thus, we may assume that *E*(*G*) *∅*.

Now, let the induction statement be true for *n < t* −1 and |*V* (*G*) | = *t*. Assume *S* consists of all the source nodes in *G* and *G*^*′*^ is obtained by removing all nodes in *S* together with their outgoing edges from *G*. Then, using Lemma 4, ℬ (*G*^*′*^, *k*) is uniquely constructed from ℬ (*G, k*) as the source nodes are uniquely determined. Thus, the induction assumption on *n* implies that *E*(*G*^*′*^) and *V*_*l*_(*G*^*′*^) are uniquely determined for any 0 *≤ l ≤* depth(*G*^*′*^).

In addition, since *V*_*l*_(*G*) = *V*_*l−*1_(*G*^*′*^) for 1 ≤*l* ≤ depth(*G*) and *S* = *V*_0_(*G*) are uniquely determined, *V*_*l*_(*G*) is uniquely determined for 0≤ *l* ≤ depth(*G*).

Now, we show that each edge (*u, v*) ∈ *E*(*G*) with *u* ∈ *S* is uniquely determined.

If *k* = 1, (*u, v*) ∈ *E*(*G*) if and only if *v* ∈ *B*_*G,k*_(*u*). Therefore, (*u, v*) is uniquely determined. Let *k* ≥2. Then, we determine each edge (*u, v*) with *u*∈*S* recursively as follows.

If *v* ∈*V*_1_(*G*), there is no directed path of length greater than one from *u* to *v*. Therefore, (*u, v*) ∈ *E*(*G*) if and only if *v* ∈ *B*_*G,k*_(*u*). This implies that (*u, v*) is uniquely determined.

Now, assume we have determined each (*u, v*) ∈ *E*(*G*) with *u*∈*S* and *v* ∈*V*_*l*_(*G*) for 1 ≤ *l* ≤ *m* −1. We then aim to determine each edge (*u, v*) with *v* ∈ *V*_*m*_(*G*) and *u* ∈ *S*. To do so, let *P* : *v*_0_, *v*_1_, …, *v*_*b*_ be a directed path of length 2 ≤ *b* ≤ *k*, where *v*_0_ = *u, v*_*b*_ = *v*, and *v*_1_ ≠ *u, v*. Then, by Observation 3, *v*_1_ ∈ *V*_*l*_(*G*), where *l < m*. Thus, the edge (*v*_0_, *v*_1_) is uniquely determined. Additionally, for 1 ≤ *i* ≤ *b* − 1, (*v*_*i*_, *v*_*i*+1_) ∈ *E*(*G*^*′*^) is uniquely determined by the induction assumption on *n*. Thus, *P* is uniquely determined, implying that the number of directed paths of length 2 ≤ *k* from *u* to *v* is unique to *B*(*G, k*). Now, assume that *n* (*u, v*) is the number of such paths. Then, *n*(*u, v*) −*n*^*′*^(*u, v*) uniquely determines the edge (*u, v*).

Therefore, *E*(*G*) = *E*(*G*^*′*^) ∪ {(*u, v*) ∈ *E*(*G*) : *u* ∈ *S*} is uniquely determined, which concludes the proof.

#### Proposition 4.

*Let G and* 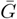 *be two labeled DAGs. If k* ≥ max {*depth*(*G*), *depth* 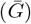}, *then the s-k-RF distance between G and* 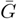 *equals the s-k*^*′*^*-RF distance between G and* 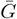 *for any k*^*′*^ *> k*.

**Proof**. If *k* ≥ max{depth(*G*), depth 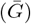}, then *D*_*X,k*_(*v*) = *D*_*X*_ (*v*) *∪* {*v*} for *X* = *G*, 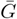 and *v* ∈ *V* (*X*). Thus, 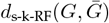 is independent of *k*, implying that 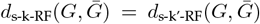 is the same for any *k*^*′*^ *> k*.

#### Proposition 5.

*Let G and* 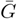 *be two labeled DAGs and let* 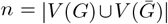 *and* 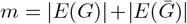. *The time complexity to compute* 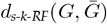 *is O*(*kn*(*n* + *m*)).

**Proof**. The proof appears in Section II, Supplementary Document.

#### Example 2.

*Consider the DAGs in Figure 8. For k* = 1, 2, 3 *and* 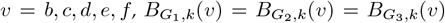 *are listed below*.

**Fig. 8.**
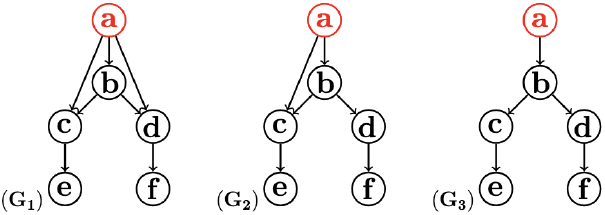
Three rooted labeled DAGs *G*_1_, *G*_2_, and *G*_3_. For each *k* = 1, 2, 3, the pairwise s-*k*-RF between them is equal to 2.

| $v \backslash k$ | 1 | 2 | 3 |
| --- | --- | --- | --- |
| $b$ | $\{b, c, d\}$ | $\{b, c, d, e, f\}$ | $\{b, c, d, e, f\}$ |
| $c$ | $\{c, e\}$ | $\{c, e\}$ | $\{c, e\}$ |
| $d$ | $\{d, f\}$ | $\{d, f\}$ | $\{d, f\}$ |
| $e$ | $\{e\}$ | $\{e\}$ | $\{e\}$ |
| $f$ | $\{f\}$ | $\{f\}$ | $\{f\}$ |

*In addition, B*_*X,k*_(*a*) *for X* = *G*_1_, *G*_2_, *G*_3_ *and k* = 1, 2, 3 *have the following values*.

In addition, $B_{X,k}(a)$ for $X = G_1, G_2, G_3$ and $k = 1, 2, 3$ have the following values.
| $X \backslash k$ | 1 | 2 | 3 |
| --- | --- | --- | --- |
| $G_1$ | $\{a, b, c, d\}$ | $\{a, b, c^2, d^2, e, f\}$ | $\{a, b, c^2, d^2, e, f\}$ |
| $G_2$ | $\{a, b, c\}$ | $\{a, b, c^2, d, e\}$ | $\{a, b, c^2, d, e, f\}$ |
| $G_3$ | $\{a, b\}$ | $\{a, b, c, d\}$ | $\{a, b, c, d, e, f\}$ |

*Consequently, for k* = 1, 2, 3,

*d*_*s-k-RF*_(*G*_1_,*G*_2_) = *d*_*s-k-RF*_(*G*_1_,*G*_3_) = *d*_*is-k-RF*_(*G*_2_,*G*_3_) = 2.

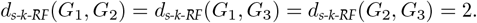

Example 2 shows that the s-*k*-RF measures cannot naively capture the dissimilarity of the labeled DAGs, as we expect *d*_s-k-RF_(*G*_1_, *G*_2_) *< d*_s-k-RF_(*G*_1_, *G*_3_). In fact, when the number of redundant edges incident to the same source node in a DAG (e.g., *G*_1_) varies and this is the only change to the DAG, the distance of the new DAG (e.g., *G*_2_ or *G*_3_) from the old one is always a fixed number regardless of how many edges are removed from or added to the DAG. However, the number of added or removed edges affects the number of edges. Thus, the measures are not efficiently sensitive to the changes occurring in the number of edges. This caveat motivates us to define a modified version of the measures.

## 5 M-*k*-RF Distances for labeled DAGs

Let *G* be a labeled DAG, and *k* ≥ 0 be an integer. We define:

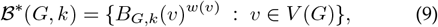

where *w*(*v*) = 1 if *v* is a root, and *w*(*v*) = indegree(*v*) otherwise.

### Definition 5.

*Let G and* 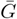 *be two labeled DAGs*.*The modified*

*k-RF (m-k-RF) distance between G and* 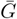 *is defined as follows:*

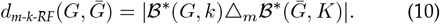

### Example 3.

*Consider DAGs in Figure 8. Then, for each k* = 1, 2, 3, ℬ^∗^(*G*_1_, *k*), ℬ^∗^(*G*_2_, *k*), *and* ℬ^∗^(*G*_3_, *k*) *are respectively:*

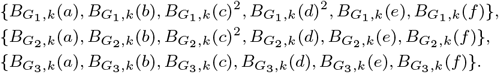

*Thus, for any k* = 1, 2, 3, *we have:*

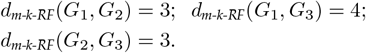

### 5.1 Mathematical Properties

#### Lemma 5.

*For three multisets A, B, and C, the following inequality holds:*

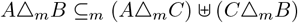

**Proof**. If *x*^*m*(*x*)^ *∈ AΔ*_*m*_*B*, then *x*^*m*(*x*)^ *∈ A \*_*m*_ *B* or *x*^*m*(*x*)^ *∈ B \*_*m*_ *A*. Suppose *x*^*m*(*x*)^ *∈ A\*_*m*_ *B*. Then, *m*(*x, A*) *> m*(*x, B*). If *x ∈/* Supp(*C \*_*m*_ *B*), we have *m*(*x, A*) *> m*(*x, B*) ≥ *m*(*x, C*). This implies that *x ∈* Supp(*A \*_*m*_ *C*) and

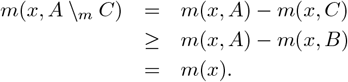

Thus, *m*(*x*) ≤ *m*(*x, A* Δ_*m*_*C*) + *m*(*x, C* Δ_*m*_*B*).

On the other hand, if *x* ∈ Supp(*C \*_*m*_ *B*) and *m*(*x, C*) ≥ *m*(*x, A*), we have:

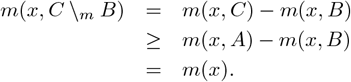

If *x* ∈ Supp(*C \*_*m*_ *B*) and *m*(*x, C*) *< m*(*x, A*), we have *m*(*x, A*) *> m*(*x, C*) *> m*(*x, B*), implying *x* ∈ Supp(*A \*_*m*_ *C*). Thus, we have:

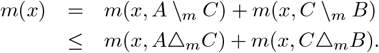

Lastly, if *x*^*m*(*x*)^ ∈ *B \*_*m*_ *A*, we can obtain the same result. In summary, we have:

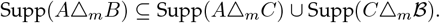

In addition, for each *x ∈* Supp(*AΔ*_*m*_*B*), we have:

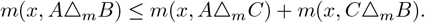

Thus, the inequality holds.

#### Lemma 6.

*Let k* ≥ 0 *be an integer. The m-k-RF measure satisfies the non-negativity, symmetry, and triangle inequality conditions*.

**Proof**. The non-negativity and symmetry conditions follow from the definition of the measures. For the triangle inequality, let *G*_1_, *G*_2_, and *G*_3_ be three labeled DAGs, and let *A* = ℬ^∗^(*G*_1_, *k*), *B* = ℬ^∗^(*G*_2_, *k*), and *C* = ℬ^∗^(*G*_3_, *k*). We have

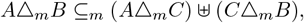

by Lemma 5. This implies that

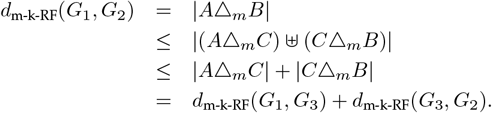

Consequently, we have

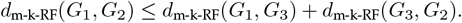

#### Theorem 6.

*Let k* ≥ 1 *be an integer. The m-k-RF measure is a metric in the space of labeled DAGs*.

**Proof** Let *G* and 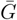 be two labeled DAGs. By Lemma 6, it is enough to show that if 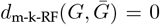, then *G* and 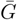 are identical. As 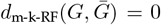 implies that 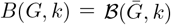, by the proof of Theorem 3, *G* and 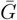 are identical.

#### Proposition 7.

*Let G and* 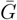 *be two DAGs. The time complexity to compute the modified k-RF distance between G and* 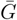 *is O*(*kn*(*n* + *m*)), *where* 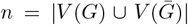 *and* 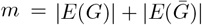

*Proof*. First, note that computing indegree of nodes in *G* and 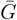 can be done in 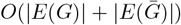 time. Therefore, the time complexity to compute the symmetric difference of ℬ ^∗^(*G, k*) and 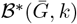 is the same as the time complexity to compute the symmetric difference of ℬ (*G, k*) and 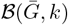. Therefore, the proof follows from the proof of Proposition 5.

### 5.2 M-*k*-RF Distance refines S-*k*-RF Distance

To demonstrate that the m-*k*-RF distance refines the s-*k*-RF distances, we used a dataset consisting of 61 randomly generated labeled DAGs with 50 nodes. For *k* = 1, 2, 3, 4, we computed the normalized s-*k*-RF and normalized m-*k*-RF distance between one DAG and every other DAG in the dataset, where the normalized s-*k*-RF and normalized m-*k*-RF distance between two DAGs *G* and 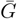 are defined as:

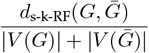

and

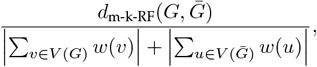

respectively. Figure 9 presents the distance values obtained by each measure between the reference DAG and every other DAG.

**Fig. 9.**
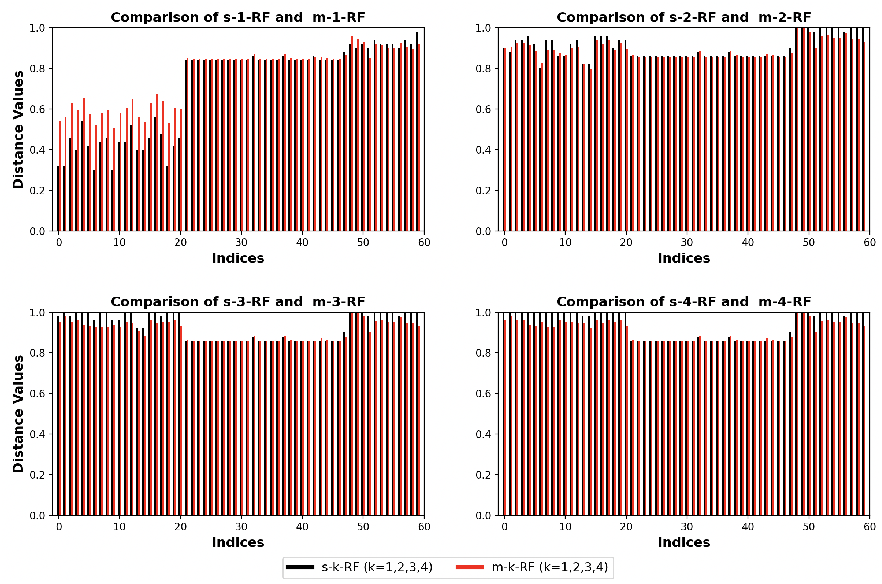
Comparison between s-*k*-RF and m-*k*-RF distances between a fixed labeled DAG *G* and every other DAG in a data set with 60 random labeled DAGs *G*_0_, *G*_1_,…,*G*_59_ for *k* = 1, 2, 3, 4. Each black (red) bar on index *i* presents *d*_s-k-RF_(*G, G*_*i*_) (*d*_m-k-RF_(*G, G*_*i*_)), where *k* = 1, 2, 3, 4.

This figure indicates that the variability between the m-*k*-RF distance values is higher than between the s-*k*-RF distance values, revealing the fact that the m-*k*-RF distances could enhance the resolution of the s-*k*-RF distances.

### 5.3 Correlation between S-*k*-RF and M-*k*-RF Distance

To determine how the s-*k*-RF distance and m-*k*-RF distance are related to each other, we created two datasets, containing 250 randomly generated labeled DAGs with 50 nodes each. DAGs in the first dataset had the same number of edges, whereas DAGs in the second dataset had different number of edges. We then computed the pairwise normalized s-*k*-RF and m-*k*-RF (1 ≤ *k* ≤ 7) distances between the DAGs in each dataset. The Pearson correlation between s-*k*-RF and m-*k*-RF distances are listed in Table 1.

**TABLE 1.** The Pearson correlation between s-*k*-RF and m-*k*-RF distances for 1 *≤ k ≤* 7.

| $k$ | Dataset 1 | Dataset 2 |
| --- | --- | --- |
| 1 | 0.93213 | 0.20302 |
| 2 | 0.94762 | 0.18250 |
| 3 | 0.94958 | 0.17914 |
| 4 | 0.94971 | 0.17865 |
| 5 | 0.94972 | 0.17855 |
| 6 | 0.94972 | 0.17852 |
| 7 | 0.94972 | 0.17852 |

The correlation analysis indicates that the s-*k*-RF and m-*k*-RF distance were positively correlated. However, the correlation coefficients for the first dataset were significantly higher than those for the second dataset. This is expected, as the m-*k*-RF distance more effectively captures dissimilarity caused by variability in the number of edges between DAGs. Additionally, the analysis shows that when *k* increased, the correlation coefficient in the first dataset increased slightly, whereas in the second dataset, it decreased with increasing *k*.

## 6 Comparison of S-*k*-RF and M-*k*-RF Dis- tances with Other DAG Distances

In this section, we compare the two *k*-RF distances with the Generalized Kendall-tau (GKT) distance [23] and Katz dissimilarity [26] (Section III, Supplementary Document).

### 6.1 Correlation of S-*k*-RF and M-*k*-RF with GKT and Katz

To assess how the s-*k*-RF distance or the m-*k*-RF distance correlates with the GKT distance and Katz dissimilarity, we computed the pairwise Katz dissimilarity and GKT distance between the DAGs in the second dataset used in Section 5.3. (A similar analysis on the first dataset yielded comparable results and is presented in Section V, Supplementary Document.)

The Katz dissimilarity and GKT distance are both parameterized measures. We computed the Katz dissimilarity scores using parameters *p* = 0.5 and *q* = 0 or 0.3, and the GKT distances using parameters *α* = 0.8 or 0.4 and *γ* = 0.0004. The Pearson correlation coefficients between the s-*k*-RF distance or the m-*k*-RF distance and the GKT or Katz measures are shown in Figure 10 for each 1 ≤ *k* ≤ 7. The analyses suggest the following:

**Fig. 10.**
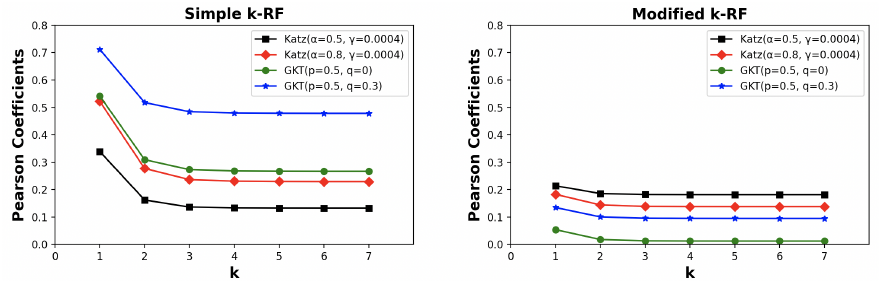
The Pearson correlation coefficients of s-*k*-RF (left) and m-*k*-RF (right) with the Generalized Kandall-tau distances and Katz dissimilarity under two parameter settings each. Each point (*k, y*) on a curve represents a correlation coefficient *y* between the corresponding GKT or Katz distance and the s-*k*-RF or m-*k*-RF distance.

a. All correlations were positive.
b. The correlation between the s-*k*-RF distance and the GKT distance was stronger than that with the Katz dissimilarity.
c. The correlation between the m-*k*-RF distance and the Katz dissimilarity was stronger than that with the GKT distances.
d. Both the GKT and Katz distances were more strongly correlated with the s-1-RF and m-1-RF distances than with the s-*k*-RF and m-*k*-RF distances for *k >* 1.

Point (d) can be attributed to the fact that both distances are based on the dissimilarity between pairs of nodes across DAGs, which aligns more closely with the s-1-RF and m-1-RF distances as they capture dissimilarity between 1-descendant-hoods across DAGs. Each 1-descendant-hood contains a node with its children, making the measures more compatible with the structure of the GKT and Katz dissimilarities.

### 6.2 Clustering DAGs Using S-*k*-RF and Other Distances

We also investigated the effectiveness of these distances in classifying DAGs. We first generated five random labeled DAGs with 50 nodes each. For each of these, we created 50 variants by deleting or reversing edges while preserving the acyclic property, resulting in five DAG families, each comprising 51 DAGs. The depth of the generated DAGs ranged from 3 to 8. Note that by Proposition 4, if *k >* 8, the s-*k*-RF score for each pair of DAGs equals its s-8-RF score.

Next, we computed the s-*k*-RF and m-*k*-RF distances (1≤ *k* ≤ 8), the GKT distance (with parameters *p* = 0.5 and *q* = 0 or 0.3), and the Katz dissimilarity (with parameters *α* = 0.5, 0.8, and *γ* = 0.0004) for every pair of DAGs within and across the five DAG families. We then applied the *K*-Means clustering algorithm to group the DAGs into *n* clusters, for *n* = 2, …, 49.

The Silhouette scores [18] of clustering the DAGs into *n* groups for *n* = 2, …, 49 for the resulting clustering are presented in Figure 11. Our findings include:

**Fig. 11.**
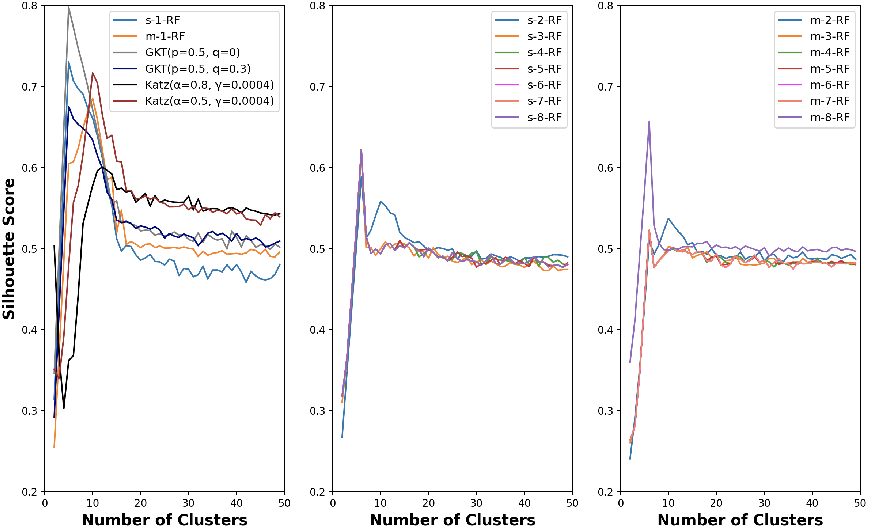
The Silhouette scores of *K*-means clustering of the labeled DAGs from five families, generated from 5 base DAGs, into *n* (2 ≤ *n* ≤ 49) groups under the s-*k*-RF distances (1 ≤ *k* ≤ 8), the GKT distance [23] (with parameters *p* = 0.5 and *q* = 0, or 0.3 and the Katz dissimilarity (wiht parameters *α* = 0.5, or 0.8 and *γ* = 0.0004).

1. Among the s-*k*-RF and m-*k*-RF distances, only s-1-RF distance accurately determined the true number of DAG families. This is evidenced by the highest Silhouette score at *n* = 5 for s-1-RF distance, a result not observed for other s-*k*-RF measures or any m-*k*-RF measures.
2. The GKT distance was another one that successfully identified the correct number.
3. The clustering performance of the s-1-RF distance is more similar to the GKT distance. This aligns with the observation we made in our correlation analysis in Section 6.1, where the correlation of the s-1-RF distance with the GKT was higher than with the other measure.
4. Although the m-*k*-RF distances indicated a stronger correlation with the GKT distance when the underlying DAGs had the same number of edges (Tables S1–S4, Section V, Supplementary Document), their clustering performance was similar to the Katz dissimilarity. This could be attributed to the existence of DAGs with different numbers of edges in the dataset. Note that according to the result from Section 6.1, the distances indicated higher correlation with Katz dissimilarity when the underlying DAGs had different numbers of edges.

## 7 Comparison of S-*k*-RF Distances and *k*-RF Distances on Trees with Different Label Sets

Recall that the s-*k*-RF distance is a simplified version of the *k*-RF distance, customized for labeled DAGs. We compare the performance of these two distances in the space of labeled trees with different label sets.

### 7.1 Clustering Trees Based on Their Label Sets

We first investigate the ability of the s-*k*-RF and *k*-RF distances to differentiate between multi-labeled tree families with different label sets. (The definitions of the measures extend naturally to multi-labeled DAGs and are presented in Section VI, Supplementary Document.)

For this purpose, we computed pairwise *k*-RF and s-*k*-RF distances between multi-labeled trees from five families, generated using the following procedure:

- 20,000 rooted labeled trees were first generated using the method described in Section 3.4. Each tree contained 30 nodes, uniquely labeled with integers from 0 to 29 (i.e., the label set is {0, 1, …, 29}).
- The generated trees were then converted into multilabeled trees using the approach described in [16]. This process involves deleting a few nodes and reattaching their children to the deleted nodes’ parents. Additionally, some edges are contracted, and the label sets of the two endpoints of each contracted edge are merged and assigned to the new node.

We then sampled 240 multi-labeled trees with the same label set of size 30. We grouped them into 5 families, each composed of 48 trees. Finally, we changed some labels so that trees from different families had label sets differing by only one mutation. The diameter and depth of each tree were at most 20 and 19, respectively. Therefore, for *k >* 19, the s-*k*-RF and *k*-RF score for each pair of trees equal its s-19-RF and 19-RF score, respectively (see Proposition 4 and Proposition 1 in [19]).

The silhouette scores for clustering trees using distance values obtained by each *k*-RF and each s-*k*-RF are presented in Figure 12 for *k* = 1, 2, …, 19.

**Fig. 12.**
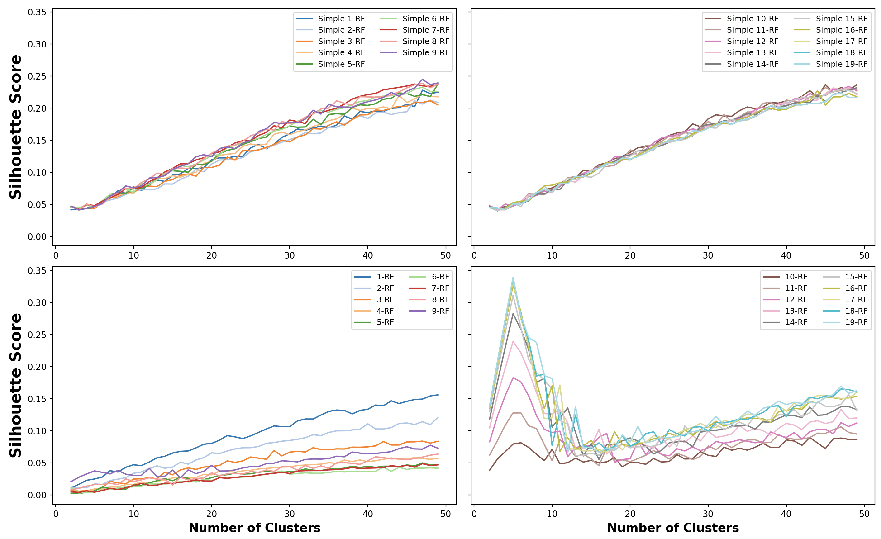
Silhouette scores of clustering 240 multi-labeled trees from five families into different number of groups under the s-*k*-RF and *k*-RF distance for *k* = 1, 2, …, 19. Trees from the same family shared the same label set, while trees from different families had label sets differing by only one label.

As Figure 12 shows, neither of the s-*k*-RF measures could recognize the number of tree families. Furthermore, the clustering performance of the measures did not change noticeably when *k* increased from 1 to 19. On the other hand, the *k*-RF distances could recognize the number of clusters for *k* ≥ 12; additionally, the clustering performance improved greatly when *k* increased from 12 to 19. This aligns with the result in Section 3.5, where we observed that the Pearson correlation between the measures is low for *k* ≥ 12 when the underlying trees had different label sets.

To see why the *k*-RF and the s-*k*-RF measures behave differently in the clustering experiment, we recall that for large *k*, the *k*-RF measure behaves similarly to the RF measure proposed in [19], which does not capture any similarity for trees with different label sets (Remark 1, [19]). This in turn facilitates differentiating between such trees. On the other hand, for large *k*, the s-*k*-RF distances become the RF distance (Definition 3 and Proposition 1) which can capture similarity between trees with different label sets.

### 7.2 Resolution of S-*k*-RF and *k*-RF Distances on Trees with Different Label Sets

Although for large *k* (*k* ≥ 12), the *k*-RF measure could cluster trees based on their label sets better than the s-*k*-RF distance, as discussed above, they cannot capture similarity of trees with different label sets well. This motivated us to compare the resolution of the *k*-RF measure and s-*k*-RF measure on trees with different label sets.

For this purpose, we computed the normalized s-*k*-RF and normalized *k*-RF distance (1 ≤ *k* ≤ 4) between one reference labeled tree and any other labeled tree indexed by 0, 1, …, 49, where the normalized s-*k*-RF and normalized *k*-RF distance between two trees *T* and 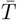 are respectively calculated as 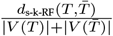 and 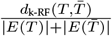.

The set of trees was a subset of the dataset used in Section 3.5; additionally, all trees except the reference tree shared the same label set.

Figure 13 shows the distance values obtained by each measure between the reference tree and any other tree indexed by 0, 1, …, 49. The figure suggests:

**Fig. 13.**
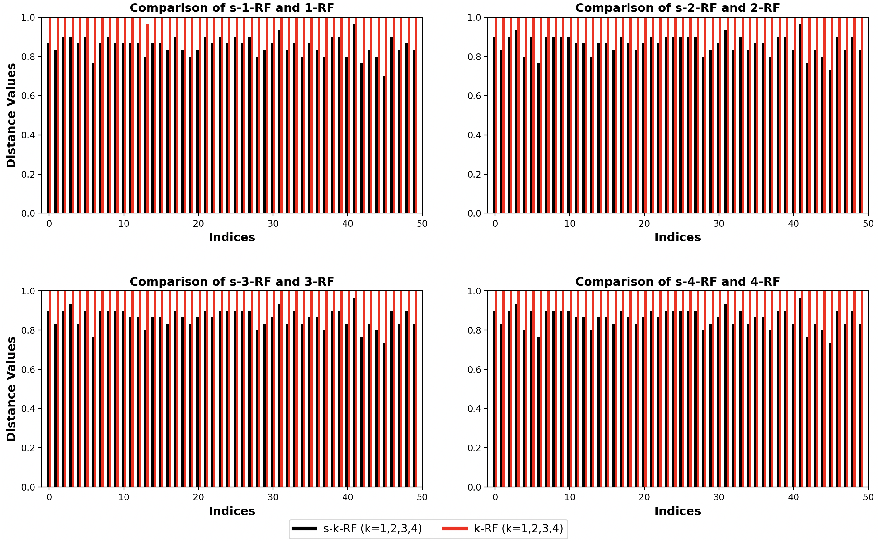
Comparison between s-*k*-RF and *k*-RF distances of a reference labeled tree with any other labeled tree indexed by 0, 1, …, 49. Each black (red) bar on index *i* presents the s-*k*-RF (*k*-RF) distance between the reference tree and the tree indexed by *i*. The label set of the reference tree was different from any other tree, and *k* ranged from 1 to 4.

- The *k*-RF measure could not capture any similarity between trees in most pairs, while the s-*k*-RF could capture similarity between any two evaluated trees.
- The variability between the distance values obtained by the s-*k*-RF measures was much higher. This indicates that the resolution of the s-*k*-RF measures for trees with different label sets seems to be higher, compared to the *k*-RF distances.

## 8 Discussions and Conclusions

We proposed the s-*k*-RF distance for labeled rooted trees by refining the *k*-RF measures introduced by [19]. We further extended the s-*k*-RF distances to m-*k*-RF distances for DAGs. Note that both the s-*k*-RF and m-*k*-RF distances have been generalized to the space of multi-labeled DAGs (Section VI, Supplementary Document).

Our simulation experiments indicate that the distribution of the pairwise s-*k*-RF distances is approximately normal in the space of rooted labeled trees with *n* nodes, for small *n* and each *k* ≤ *n* −1, This behavior differs from that observed for the *k*-RF measure in [19].

We also conducted a correlation analysis by computing the Pearson correlation between s-*k*-RF measures (1≤ *k* ≤ 21) and the three existing measures CASet ∩, DISC ∩, and GRF [11], [22], which are developed for comparison of cancer mutation trees. For each of the three measures, we observed that the correlation remained nearly constant across different values of *k* ≥ 1, in contrast to the findings of [19], which reported noticeable fluctuations as *k* increased (see Figure 10 in [19]).

We compared the *k*-RF and the s-*k*-RF distance (1≤ *k* ≤ 21) using a Pearson correlation. We observed that for large *k*, the two measures were highly correlated when the underlying trees had the same label set. However, they did not show strong correlation when the trees had different label sets.

The s-*k*-RF and m-*k*-RF measures are each a metric in the space of labeled DAGs. The normalized m-*k*-RF measure seems to have higher resolution than the normalized s-*k*-RF measure on the space of labeled DAGs with the same label set. The s-*k*-RF and m-*k*-RF measures (1≤ *k* ≤ 7) showed stronger correlation when the underlying DAGs had the same number of edges, compared to the scenario in which the DAGs had different numbers of edges.

The s-*k*-RF distance (1 ≤ *k* ≤ 7) showed weaker correlation with the Katz dissimilarity [26] than with the GKT distance [23]. A similar pattern was observed for the m-*k*-RF distance when the underlying DAGs had the same number of edges. However, when the DAGs differed in the number of edges, the m-*k*-RF distance exhibited stronger correlation with the Katz dissimilarity than with the GKT distance.

Additionally, we demonstrated that the s-1-RF distance can effectively identify the number of clusters in a DAG dataset, outperforming the Katz similarity [26] and achieving comparable performance to the GKT distance [23].

The s-*k*-RF measures can efficiently compare two labeled DAGs *G* and 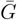 with different node sets; however, in order to compute the distance between such DAGs by other DAG measures, one needs to first make their node sets the same. In particular, to calculate the GKT distance between the DAGs, first some isolated nodes are added to the DAGs so that they obtain the same node set 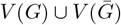 [23]. This approach can impose a high degree of dissimilarity between DAGs. This is because it induces 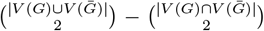 node pairs, each with a tail and/or a head in 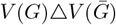. Clearly, if (*u, v*) is such a pair, 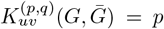 if *u* and *v* are connected by an edge in *G* or 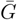 and 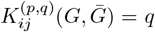 otherwise. Therefore, this strategy could overestimate the contribution of differing nodes to the distance between the DAGs. Moreover, the authors in [26] prefer to remove certain nodes from the DAGs to equalize their node sets to 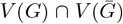. However, this approach may underestimate the contribution of differing nodes to the distance between the DAGs.

The *k*-RF measures could not capture similarity of labeled trees with different label sets well, showing a less resolution against the s-*k*-RF distances when comparing such trees. As such, the s-*k*-RF distances are more efficient in the scenarios where different labels should not contribute greatly to the distance between labeled DAGs.

We proposed an algorithm to compute the m-*k*-RF and s-*k*-RF distances between two labeled DAGs. The algorithm runs in polynomial time; however, its time complexity is not directly comparable to the algorithms in [8] and [1], which are used to calculate the RF distance between phylogenetic trees and phylogenetic networks, respectively. This caveat could be justified because our algorithm has to account for distinct labels on internal nodes, whereas phylogenetic trees or networks assign the same label to all internal nodes. Furthermore, the algorithm provides greater flexibility by allowing *k* to vary.

Future work includes investigating the application of the s-*k*-RF distances in comparison of cancer trees, cell trees and designs of machine learning algorithms. For example, Wang et al. [33] introduce a kernel to capture similarity between two DAGs *G* and 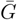 for action recognition tasks.

Each DAG corresponds to a video clip, where a node is a component consisting of the video trajectories corresponding to a moving body part, and there is a directed edge from component *c*_1_ to component *c*_2_ if the body part representing *c*_1_ moves before the one representing *c*_2_. Now, one may use the Gaussian kernel empowered by the s-*k*-RF distance. Such a kernel could be defined as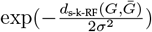, where *σ* controls the sensitivity to the distance.

Furthermore, validating these measures in terms of their contribution to DAG learning methods through techniques such as bootstrap aggregation is also planned. For example, the authors in [35] use a metric to improve their model, which learns a DAG that identifies protein biomarkers for the response to treatment in ovarian cancer. Replacing the distance with the s-*k*-RF distance could recover its potential application in biological scenarios.

In addition to the above future research plans, it is also interesting to explore the frequency distribution of the s-*k*-RF measures in the space of rooted labeled trees with a large number of nodes in line with the paper [14], which investigates the distribution of the *k*-RF measures proposed in [19].

The code to compute pairwise s-*k*-RF and m-*k*-RF distances of a set of labeled DAGs can be found in https://github.com/Elahe-khayatian/sKRF-distances.git.

## Acknowledgments

The authors thank the anonymous reviewers for their insightful and constructive suggestions on the first submission. This work was partially supported by Singapore MOE Research Grant [A-8001951-00-00].

**Elahe Khayatian** received her BSc and MSc in Mathematics. She now is a PhD candidate in the Department of Mathematics at National University of Singapore. Her research interests include Computational Biology and Graph Theory.

**Louxin Zhang** is a computational biologist. He is currently a professor in the Department of Mathematics at National University of Singapore. He is recognized for his contributions to combinatorial semigroup theory, phylogenetic trees and networks, as well spaced seeds for sequence comparison in bioinformatics.

## References

[1] T. Asano, J. Jansson, K. Sadakane, R. Uehara, and G. Valiente, “Faster Computation of the Robinson-Foulds Distance Between Phylogenetic Networks”, Inf. Sci., vol. 197, pp. 77–90, 2012.

[2] A. Basu, “Directed Acyclic Graphs to Explore Causality in Epidemiological Study Designs, part I: an introduction to DAGs”, Qeios, 3:3, 2020.

[3] G. Cardona, M. Llabrés, F. Rosselló, and G. Valiente, “Comparison of Tree-Child Phylogenetic Networks”, IEEE/ACM Trans. Comput. Biol. Bioinform., vol. 6, no. 4, pp. 552–569, 2009.

[4] G. Cardona, M. Llabrés, F. Rosselló, and G. Valiente, “Metrics for Phylogenetic Networks I: Generalizations of the Robinson-Foulds Metric”, IEEE/ACM Trans Comput Biol Bioinform, vol. 6, no. 1, pp. 46–61, 2009.

[5] G. Cardona, M. Llabrés, F. Rosselló, and G. Valiente, “Metrics for Phylogenetic Networks II: Nodal and Triplets Metrics”, IEEE/ACM Trans. Comput. Biol. Bioinform., vol. 6, no. 3, pp. 454–469, 2009.

[6] G. Cardona, M. Llabrés, F. Rosselló, and G. Valiente, “On Nakhleh’s Metric for Reduced Phylogenetic Networks”, IEEE/ACM Trans. Comput. Biol. Bioinform., vol. 6, no. 4, pp. 629–638, 2009.

[7] F. Cheng, I.A. Kovács, and A.L. Barabási, 2019. “Network-based Prediction of Drug Combinations”, Nat. Commun., 10(1), 1197, 2019.

[8] W.H.E. Day, “Optimal Algorithms for Comparing Trees with Labeled Leaves”, J. Classif., vol. 2, no. 1, pp. 7–28, 1985.

[9] R. Diaz-Uriarte, “Cancer Progression Models and Fitness Landscapes: A Many-to-Many Relationship”, Bioinform., vol. 34, no. 5, pp. 836–844, 2018.

[10] R. Diaz-Uriarte and C. Vasallo, “Every Which Way? On Predicting Tumor Evolution Using Cancer Progression Models”, PLoS Comput. Biol., vol. 15, no. 8, pp. e1007246, 2019.

[11] Z. DiNardo, K. Tomlinson, A. Ritz, and L. Oesper, “Distance Measures for Tumor Evolutionary Trees”, Bioinform., vol. 36, no. 7, pp. 2090–2097, 2020.

[12] D. Doncevic, C. Herrmann, “Biologically Informed Variational Autoencoders Allow Predictive Modeling of Genetic and Drug-Induced Perturbations”, Bioinform., vol. 39, no. 6, 2023.

[13] M. C. Fontaine, J. B. Pease, A. Steele, et al. “Extensive Introgression in A Malaria Vector Species Complex Revealed by Phylogenomics”, Science, vol. 347, no. 6217, 1258524, 2015.

[14] M. Fuchs and M. Steel, “The Asymptotic Distribution of The K-Robinson-Foulds Dissimilarity Measure on Labelled Trees”, arXiv preprint, arXiv:2412.20012, 2024.

[15] D. H. Huson, R. Rupp, and C. Scornavacca. Phylogenetic Networks. Cambridge, UK, 2012.

[16] K. Jahn, N. Beerenwinkel, and L. Zhang, “The Bourque Distances for Mutation Trees of Cancers”, Algor. Mol. Biol., vol. 16, no. 9, pp. 1–15, 2021.

[17] J. Jansson, K. Mampentzidis, R. Rajaby, and W.K. Sung, “Computing the Rooted Triplet Distance Between Phylogenetic Networks”, Algorithmica, vol. 83, pp. 1786–1828, 2021.

[18] L. Kaufman and P. Rousseeuw, Finding Groups in Data: An Introduction To Cluster Analysis. John Wiley & Sons, 2009.

[19] E. Khayatian and G. Valiente and L. Zhang, “The k-Robinson–Foulds Dissimilarity Measures for Comparison of Labeled Trees”, J. Comput. Biol., vol. 31, no. 4, pp. 328–344, 2024. A preliminery version appears in the Proceedings of the RECOMB-CG’2023, Lecture Notes in Computer Science, vol. 13883, pp. 146–161, 2023.

[20] P. Lecca, N. Casiraghi, and F. Demichelis, “Defining Order and Timing of Mutations During Cancer Progression: The TO-DAG Probabilistic Graphical Model”, Front. Genet., vol. 6, no. 309, 2015.

[21] M. Llabrés, F. Rosselló, and Gabriel Valiente, “A Generalized Robinson-Foulds Distance for Clonal Trees, Mutation Trees, and Phylogenetic Trees and Networks”, Proc. 11th ACM Int. Conf. Bioinform. Comput. Biol. Health Inform., pp. 13:1–13:10, 2020.

[22] M. Llabrés, F. Rosselló, G. Valiente, “The Generalized Robinson-Foulds Distance for Phylogenetic Trees”, J. Comput. Biol., vol. 28, no. 12, pp. 1–15, 2021.

[23] E. Malmi, N. Tatti, and A. Gionis, “Beyond Rankings: Comparing Directed Acyclic Graphs”, Data Min Knowl Disc, no. 29, pp. 1233–1257, 2015.

[24] M. M. Neyshabouri and J. Lagergren, “ToMExO: A Probabilistic Tree-Structured Model for Cancer Progression”, PLoS Comput. Biol., vol. 18, no. 12, 2022.

[25] L. Nakhleh, “A Metric on the Space of Reduced Phylogenetic Networks”, IEEE/ACM Trans. Comput. Biol. Bioinform., vol. 7, no. 2, pp. 218–222, 2010.

[26] G. Nayak, S. Dutta, D. Ajwani, P. Nicholson, and A. Sala, “Automated Assessment of Knowledge Hierarchy Evolution: Comparing Directed Acyclic Graphs”, Inform. Retr., vol. 22, pp. 256–284, 2019.

[27] J-P. Onnela, J. Saramäki, J. Hyvönen, G. Szabó, D. Lazer, K. Kaski, J. Kertész, and A-L. Barabási, “Structure and Tie Strengths in Mobile Communication Networks.” Proc. Nat. Acad. Sci., 104, no. 18 (2007): 7332–7336.

[28] D. F. Robinson and L. R. Foulds, “Comparison of Phylogenetic Trees”, Math. Biosci., vol. 53, no. 1–2, pp. 131–147, 1981.

[29] N. Rossi, N. Gigante, N. Vitacolonna, and C. Piazza, “A Conservative Approach for Describing Cancer Progression”, bioRxiv [Preprint], 2022.

[30] M. A. Steel and D. Penny, “Distributions of Tree Comparison Metrics: Some New Results”, Syst. Biol., vol. 42, no. 2, pp. 126–141, 1993.

[31] T. H. Tan, J. H. Hus, and S. H. Liu, Y. F. Huang, and M. Gochoo, “Using Direct Acyclic Graphs to Enhance Skeleton-Based Action Recognition with a Linear-Map Convolution Neural Network”, Sensors, vol. 21, no. 9, pp. 1–13, 2021.

[32] J. Wang and M. Guo, “A Metric on the Space of kth-order reduced Phylogenetic Networks”, Sci. Reports, vol. 7, pp. 1–10, 2017.

[33] L. Wang and H. Sahbi, “Directed Acyclic Graph Kernels for Action Recognition”, Proc. IEEE Int’l Confer. Comput. Vision, pp. 3168–3175, 2013.

[34] T. C. Williams, C. C. Bach, N. B. Matthiesen, T. B. Henriksen, and L. Gagliardi, “Directed Acyclic Graphs: A Tool for Causal Studies in Paediatrics”, Pediatr Res., vol. 84, no. 4, pp. 487–493, 2018.

[35] Q. Yu, S. Chowdhury, R. Wang, C.J. Huntoon, L.M. Karnitz, and S.H. Kaufmann, S.P. Gygi, M.J. Birrer, A.G. Paulovich, J. Peng, and P. Wang, “DAGBagM: Learning Directed Acyclic Graphs of Mixed Variables with an Application to Identify Protein Biomarkers for Treatment Response in Ovarian Cancer”, BMC Bioinform., vol. 23, no. 1, 2022.

[36] Y. Zinati, A. Takiddeen, and A. Emad: GRouNdGAN, “GRN-guided Simulation of Single-Cell RNA-Seq Data Using Causal Generative Adversarial Networks”, Nat. Commun., vol. 15, no. 1, 2024.

