## Supplementary Document for "Simple k-RF Metrics for Comparison of Labeled DAGs"

#### I. CASet∩, DISC∩, and GRF Distances

Let $A$ and $B$ be two sets. Then, the Jaccard distance $d_J$ between $A$ and $B$ is defined as follows:

$$d_J(A,B) = \frac{|A \triangle B|}{|A \cup B|}. \quad (11)$$

Now, let $T$ be a labeled rooted tree and $v \in V(T)$. Then, recall from [11] that $\bar{A}_T(v)$ is defined as $\bar{A}_T(v) = A_T(v) \cup \{v\}$; then, for $u, v \in V(G)$, $C_T(u,v)$ and $D_T(u,v)$ are defined as follows:

$$C_T(u,v) = \bar{A}_T(u) \cap \bar{A}_T(v), \quad D_T(u,v) = \bar{A}_T(u) \setminus \bar{A}_T(v).$$

Let $S$ and $T$ be two labeled rooted trees, $V = V(S) \cap V(T)$, and $|V| = n$. Then, CASet∩ and DISC∩ are defined as follows [11]:

$$\text{CASet} \cap (S,T) = \frac{1}{\frac{n(n-1)}{2}} \sum_{\{u,v\} \subseteq V} d_J(C_S(u,v), C_T(u,v)),$$

$$\text{DISC} \cap (S,T) = \frac{1}{n(n-1)} \sum_{\substack{(u,v) \in V \times V \\ u \neq v}} d_J(D_S(u,v), D_T(u,v)).$$

In addition, recall from [21] that clone($T$) is defined as follows:

$$\text{clone}(T) = \{\bar{A}_T(v) \ : \ v \in V(T)\}. \quad (12)$$

Next, $D_{d_J}$ between $A$ and $B$ is defined as follows [21]:

$$D_{d_J}(A,B) = \frac{\sum_{a \in A} \sum_{b \in B \setminus A} d_J(a,b)}{|A \cup B||A|} + \frac{\sum_{a \in A \setminus B} \sum_{b \in B} d_J(a,b)}{|A \cup B||B|}.$$

Finally, the Generalized Robinson-Foulds distance (GRF) between $S$ and $T$ is defined as $D_{d_J}(\text{clone}(S), \text{clone}(T))$ [21].

#### II. Proof of Proposition 5 in Section 4.1

**Lemma S1.** *Let $k \geq 0$ and $G$ be a labeled DAG. All label multi-subsets $B_{G,k}(v)$ for $v \in V(G)$ can be computed in $O(k|V(G)|(|V(G)| + |E(G)|))$ time.*

**Proof.** Take the following steps:

(a) Perform a topological sorting of $G$ in $O(|V(G)| + |E(G)|)$ time, utilizing DFS traversal.

(b) Let $V = [v_0, v_2, ..., v_{n-1}]$, where $v_i$s are arranged in the topological order, and $v \in V(G)$ is fixed. Define a $n \times (k+1)$ table $A$, where $A[i][j]$ represents the number of directed paths from $v$ to $v_i$ with length $j$. Initialize $A[i][0] = 1$ if $v_i = v$ and $A[i][j] = 0$ if $v_i \neq v$. This step can be done in $O((k+1)|V(G)|)$ time.

(c) For each $0 \leq i \leq n-1$ and each $0 \leq j \leq k-1$, if $A[i][j] > 0$, for each edge $(v_i, v_k)$, set $A[k][j+1] = A[k][j+1] + A[i][j]$. This step can be done in $O(k|E(G)|)$ time.

(d) Initialize $B_v = [\ ]$. Then, for each $0 \leq i \leq n-1$, if $v_i = v$, set $n_v = 1$; otherwise set $n_v = \sum_{j=1}^{k} A[i][j]$ and add $n_v$ to the list $B_v$. In doing so, $B_v$ represents $B_{G,k}(v)$ as a vector in which the $i$th position is $n(v, v_i)$. This step can be done in $O(k|V(G)|)$ time.

Therefore, computing all label multi-subsets $B_{G,k}(v)$ for $v \in V(G)$ can be done in $O(k|V(G)|(|V(G)| + |E(G)|))$ time. □

**Proof of Proposition 5** Let $G$ and $\bar{G}$ be two labeled DAGs and $V(G) \cup V(\bar{G}) = \{v_0, v_1, ..., v_{n-1}\}$. Now, take the following steps:

(a) Compute $B_{X,k}(v)$ for all $x \in V(X)$ and $X = G, \bar{G}$, in $O(k(|V(G)|(|V(G)| + |E(G)|) + |V(\bar{G})|(|V(\bar{G})| + |E(\bar{G})|)))$ time (see Lemma S1).

(b) For $X = G, \bar{G}$ and $x \in V(X)$, represent $B_{X,k}(x)$ by an $n$-dimensional vector $B_{X,x}$ in which $B_{X,x}(i) = n(x, v_i)$ if $v_i \in V(X)$ and $B_{X,x}(i) = 0$ otherwise. To do this, for each $x \in V(X)$, define an array $B_x$ of length $n$. Initialize $B_x[i] = 0$ for each $0 \leq i \leq n-1$. Then, for each $v_i \in V(X)$, update $B_x[i]$ to be $n(x, v_i)$. This can be done in $O(|V(X)|(n + |V(X)|))$ time for all $x \in V(X)$.

(c) Radix-sort $B(G,k)$ and $B(\bar{G},k)$ in $O(n(|V(G)| + D(G) + |V(\bar{G})| + D(\bar{G})))$ time, where $D(X) = \max\{d_{in}(x) \ : \ x \in V(X)\}$ for $X = G, \bar{G}$.

(d) Compute the symmetric difference of $B(G,k)$ and $B(\bar{G},k)$ in $O(n(|V(G)| + |V(\bar{G})|))$ time.

Now, as $|V(X)| \leq n$ and $D(X) \leq |E(X)|$ for $X = G, \bar{G}$, the total time complexity is

$$O(nk(|V(G)| + |V(\bar{G})| + |E(G)| + |E(\bar{G})|)).$$

□

### III. The GKT Distance and Katz Similarity

Let $G$ and $\bar{G}$ be two labeled DAGs on the same vertex set $V$ and $0 \leq p, q \leq 1$. In addition, suppose that there is a fixed order on $V$. Then, recall from [23] that, for each pair of nodes $i$ and $j$, $K_{ij}^{(p,q)}(G, \bar{G})$ is defined as follows:

(a) If either $(i, j) \in E(G) \cap E(\bar{G})$ or $(j, i) \in E(G) \cap E(\bar{G})$, $K_{ij}^{(p,q)}(G, \bar{G}) = 0$;
(b) If either $(i, j) \in E(G)$ but $(j, i) \in E(\bar{G}))$ or $(j, i) \in E(G)$ and $(i, j) \in E(\bar{G})$, $K_{ij}^{(p,q)}(G, \bar{G}) = 1$;
(c) If either $\{(i, j), (j, i)\} \cap E(G) \neq \emptyset$ but $\{(i, j), (j, i)\} \cap E(\bar{G}) = \emptyset$ or $\{(i, j), (j, i)\} \cap E(G) = \emptyset$ but $\{(i, j), (j, i)\} \cap E(\bar{G}) \neq \emptyset$, $K_{ij}^{(p,q)}(G, \bar{G}) = p$;
(d) If both $\{(i, j), (j, i)\} \cap E(G) = \emptyset$ and $\{(i, j), (j, i)\} \cap E(\bar{G}) = \emptyset$, then $K_{ij}^{(p,q)}(G, \bar{G}) = q$.

Then, the GKT distance $K^{(p,q)}(G, \bar{G})$ is defined as follows [23]:

$$K^{(p,q)}(G, \bar{G}) = \sum_{(i,j) \in V \times V : i < j} K_{ij}^{(p,q)}(G, \bar{G}). \tag{13}$$

Let $G$ be a DAG and $u, v \in V(G) = \{v_1, v_2, ..., v_n\}$. In addition, suppose that $0 < \alpha < 1$ is a fixed parameter. Recall from [26] that Katz Similarity (KS) between $u$ and $v$ is defined as follows:

$$KS_G(u, v) = \sum_l n_l \alpha^l, \tag{14}$$

where $n_l$ is the number of directed paths with length $l$ from $u$ to $v$. Now, let $m_{ij} = KS_G(v_i, v_j)$. The KSV for $G$ is defined as follows [26]:

$$KSV_G = (m_{11}, \cdots, m_{1n}, \cdots, m_{n1}, \cdots m_{nn}). \tag{15}$$

Let $G$ and $\bar{G}$ be two DAGs on the same vertex set $V$. Then, the KGS [26] between $G$ and $\bar{G}$ is defined as follows:

$$KGS(G, \bar{G}) = \frac{2}{1 + exp(\gamma.||KSV_G - KSV_{\bar{G}}||_1)}, \tag{16}$$

where $\gamma > 0$ is a tunable parameter which controls the sensitivity of the measure and $||.||_1$ is the $L_1$-norm.

Note that, as discussed by the authors in [26], $\gamma$ is a normalization factor and is usually set to $\frac{1}{|V||V|}$.

Furthermore, we define the Katz Graph Dissimilarity (KGD) between $G$ and $\bar{G}$ as follows:

$$KGD(G, \bar{G}) = 1 - KGS(G, \bar{G}). \tag{17}$$

### IV. Supplementary Figures for Section 3.3

We investigated the frequency distribution of the s-$k$-RF measures for rooted labeled $n$-node trees with the same label set, where $n = 7, 10, 13$ and $k = 1, \ldots, n-1$. For $n = 7$, we computed the pairwise s-$k$-RF scores for all trees in the space of rooted labeled $n$-node trees.

For $n = 10$ and $n = 13$, since the number of rooted labeled $n$-node trees is large, we computed the pairwise s-$k$-RF scores for trees in a random subset containing $n \times 1000$ rooted labeled $n$-node trees. As shown in Figures S1, S2, and S3, the distributions appear to be normal, with each red line representing a Gaussian function.

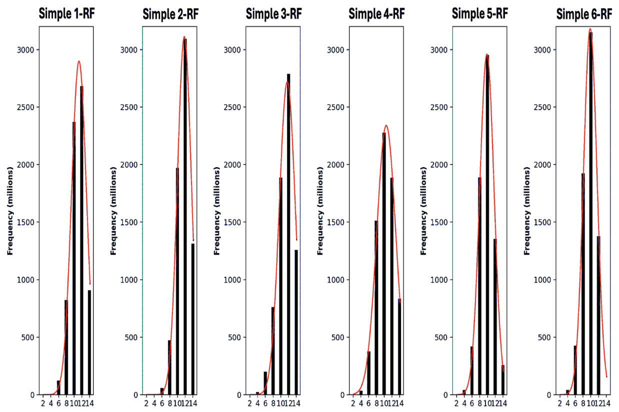


Figure S1. The distribution of s-$k$-RF distances in the space of rooted labeled 7-node trees with the same label set for $k = 1, 2, 3, 4, 5, 6$. In each panel, the red line represents a Gaussian function fitted to the bar chart using the `curve_fit` function from the SciPy library. The Gaussian function is defined as $y = a \cdot \exp\left(\frac{-(x-b)^2}{2c^2}\right)$.

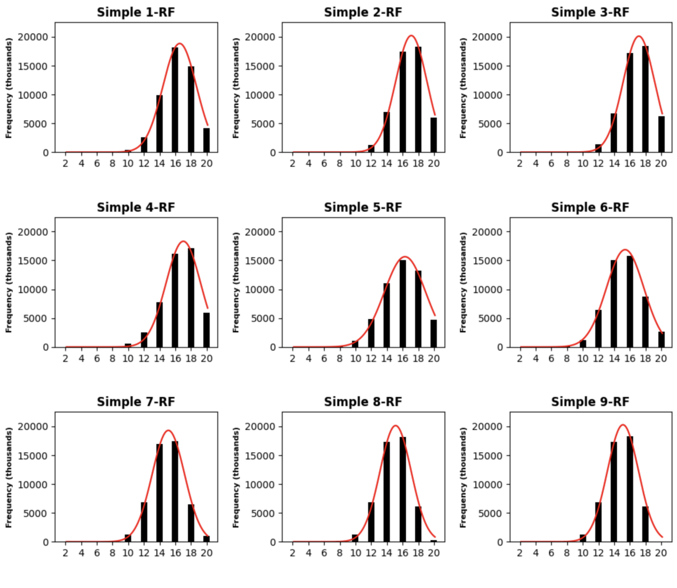


Figure S2. An overview of the distribution of s-$k$-RF scores for $k = 1, ..., 9$ in the space of 10000 randomly sampled rooted labeled 10-node trees with the same label set. In each panel, the red line represents a Gaussian function fitted to the bar chart using the `curve_fit` function from the SciPy library. The Gaussian function is defined as $y = a \cdot \exp\left(\frac{-(x-b)^2}{2c^2}\right)$.

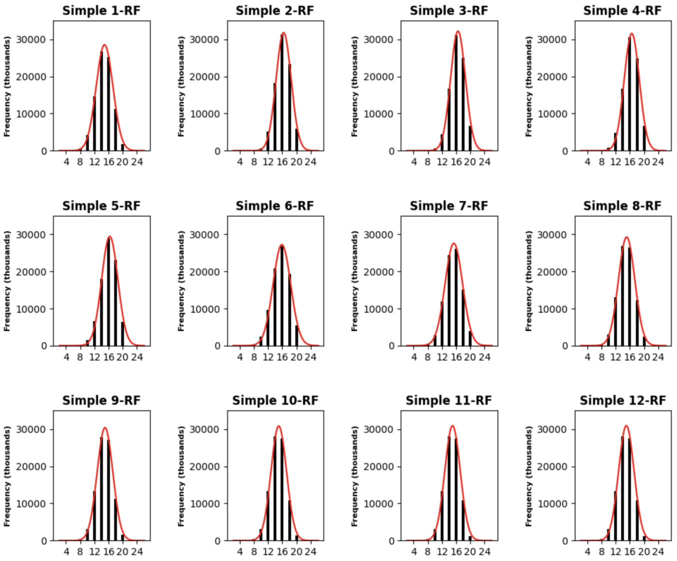


Figure S3. An overview of the distribution of s-$k$-RF scores for $k = 1, ..., 12$ in the space of 13000 randomly sampled rooted labeled 13-node trees with the same label set. In each panel, the red line represents a Gaussian function fitted to the bar chart using the `curve_fit` function from the SciPy library. The Gaussian function is defined as $y = a \cdot \exp\left(\frac{-(x-b)^2}{2c^2}\right)$.

#### V. Supplementary Tables for Section 6.1

We performed a correlation analysis similar to the analysis described in Section 6.1 utilizing the first dataset used in Section 5.3.

The Pearson coefficients of the s-$k$-RF with other DAG distances are presented in Tables S1 and S2. In addition, the Pearson coefficients of the m-$k$-RF with other DAG distances are presented in Tables S3 and S4.

TABLE S1
The Pearson correlation coefficients between the s-$k$-RF distances and the Katz dissimilarity, where $k = 1, 2, .., 7$.

| $k$ | $Katz(\alpha = 0.5, \gamma = 0.0004)$ | $Katz(\alpha = 0.8, \gamma = 0.0004)$ |
|---|---|---|
| 1 | 0.65020 | 0.31087 |
| 2 | 0.66102 | 0.31728 |
| 3 | 0.65419 | 0.31103 |
| 4 | 0.65258 | 0.30920 |
| 5 | 0.65225 | 0.30887 |
| 6 | 0.65221 | 0.30883 |
| 7 | 0.65219 | 0.30880 |

TABLE S2
The Pearson correlation coefficients between the s-$k$-RF distances and the GKT distance, where $k = 1, 2, .., 7$.

| $k$ | $GKT(p = 0.5, q = 0)$ | $GKT(p = 0.5, q = 0.3)$ |
|---|---|---|
| 1 | 0.80648 | 0.80648 |
| 2 | 0.80209 | 0.80209 |
| 3 | 0.79647 | 0.79647 |
| 4 | 0.79539 | 0.79539 |
| 5 | 0.79527 | 0.79527 |
| 6 | 0.79523 | 0.79523 |
| 7 | 0.79520 | 0.79520 |

The analyses suggest:

TABLE S3
The Pearson correlation between the m-$k$-RF distances and the Katz dissimilarity, where $k = 1, 2, .., 7$.

| $k$ | $Katz(\alpha = 0.5, \gamma = 0.0004)$ | $Katz(\alpha = 0.8, \gamma = 0.0004)$ |
|---|---|---|
| 1 | 0.69334 | 0.37687 |
| 2 | 0.68255 | 0.36552 |
| 3 | 0.67403 | 0.35758 |
| 4 | 0.67255 | 0.35584 |
| 5 | 0.67229 | 0.35560 |
| 6 | 0.67224 | 0.35555 |
| 7 | 0.67223 | 0.35554 |

TABLE S4
The Pearson correlation coefficients between the m-$k$-RF distances and the GKT distance, where $k = 1, 2, .., 7$.

| $k$ | $GKT(p = 0.5, q = 0)$ | $GKT(p = 0.5, q = 0.3)$ |
|---|---|---|
| 1 | 0.77515 | 0.77515 |
| 2 | 0.76143 | 0.76143 |
| 3 | 0.75550 | 0.75550 |
| 4 | 0.75462 | 0.75462 |
| 5 | 0.75450 | 0.75450 |
| 6 | 0.75445 | 0.75445 |
| 7 | 0.75444 | 0.75444 |

(a) The correlation values were all positive.
(b) The correlation of the GKT distance with the m-$k$-RF and s-$k$-RF distances was stronger than the correlation of the Katz dissimilarity with the distances.
(c) The GKT distance was more strongly correlated with the s-1-RF (m-1-RF) distance than with other s-$k$-RF (m-$k$-RF) distances.
(d) The Katz dissimilarity was more strongly correlated with the s-2-RF (m-1-RF) than with other s-$k$-RF (m-$k$-RF) distances.

A comparison of the analysis with the one performed in Section 6.1 suggests that when the underlying DAGs had the same number of edges, each m-$k$-RF (and s-$k$-RF) distance was more correlated with the GKT distance and the Katz dissimilarity with $\alpha = 0.5$. Furthermore, in the scenario, the change in the correlation values was less noticeable when $k$ increased from 1 to 7.

#### VI. An Extension to Multi-labeled DAGs

Let $G$ be a multi-labeled DAG with label set $X$. Each $v \in V(G)$ induces a multiset as:

$$B_{G,k}(v) = \uplus_m \{\ell(u)^{n(v,u)} \ : \ u \in D_{G,k}(v)\},$$

where $D_{G,k}(v)$ is defined in the same way as in Eqn. (1) and $n(v,u)$ is the number of directed paths from $v$ to $u$.

Now, we define $B(G,k)$ as a multiset consisting of all $B_{G,k}(v)$ with $v \in V(G)$. Therefore, by replacing $\triangle$ with $\triangle_m$ in Eqn. (8), the s-$k$-RF score for a pair of multi-labeled DAGs is well defined.

In addition, $B^*(G,k)$ is defined in the same way as in Eqn. (9). In doing so, the modified simple $k$-RF score for a pair of multi-labeled DAGs is also well defined.

**Proposition S8.** *The m-$k$-RF and s-$k$-RF distances are pseudometric in the space of multi-labeled DAGs. In other words, they satisfy the non-negativity, symmetry, and triangle inequality conditions.*

*Proof.* Let $G_1$, $G_2$, and $G_3$ be three mutation trees. Then, assuming $A = \mathcal{B}(G_1,k)$, $B = \mathcal{B}(G_2,k)$, and $C = \mathcal{B}(G_3,k)$; we have

$$A \triangle_m B \subseteq_m (A \triangle_m C) \uplus (C \triangle_m B),$$

from Lemma 5. This implies that

$$\begin{aligned} d_{\text{s-k-RF}}(G_1,G_2) &= |A \triangle_m B| \\ &\leq |(A \triangle_m C) \uplus (C \triangle_m B)| \\ &\leq |A \triangle_m C| + |C \triangle_m B| \\ &= d_{\text{s-k-RF}}(G_1,G_3) + d_{\text{s-k-RF}}(G_3,G_2). \end{aligned}$$

Thus, the triangle inequality holds. In addition, the non-negativity and symmetry conditions follow from the definition of the s-$k$-RF measures. $\square$
